# The association between dietary habits and periodontal disease in young adult women

**DOI:** 10.1101/577155

**Authors:** Akemi Hosoda, Yuriko Komagamine, Manabu Kanazawa, Yohei Hama, Akiko Kojo, Shunsuke Minakuchi

## Abstract

**Background:** Among middle-aged and elderly individuals, dietary habits have previously been reported to differ between patients with and without periodontal disease. However, in young adults, there are only a few reports that show a correlation between nutrient/food intake and periodontal disease. Moreover, no report has assessed the correlation between dietary habits measured by a self-administered diet history questionnaire (DHQ) and periodontal disease. Therefore, we assessed the correlation between dietary habits, determined using a DHQ, and periodontal disease in young adult women who are likely to develop a periodontal disease.

**Methods:** The participants were 120 healthy, non-smoking, female college students (mean age, 20.4 ± 1.1 years) from two universities who did not have any systemic disease. The participants were assessed for periodontal disease according to community periodontal index (CPI) and were divided into two groups. Subjects with a CPI code of 0, 1, or 2 were assigned to non-periodontal disease group (non-PD), and subjects with a CPI code of 3 or 4 were assigned to periodontal disease group (PD). Dietary habits were assessed using a DHQ. In addition, physical status, level of difficulty in chewing food (dietary hardness), masticatory performance, and quality of life (QoL) were assessed.

**Results:** The PD group had a significantly lower nutrient intake of minerals, fat, water-soluble vitamins, and dietary fiber than the non-PD group. In terms of food groups, the PD group consumed significantly lesser amounts of green and yellow vegetables than the non-PD group. In addition, the PD group consumed significantly lesser amounts of hard foods than the non-PD group.

**Conclusion:** Young adult women with a periodontal disease had a significantly lower nutrient/food intake than young adult women without a periodontal disease.

## 1. Introduction

Periodontal disease is a chronic inflammatory disease that begins in the gingiva and spreads throughout the periodontal tissues, culminating to its destruction [1]. Periodontal disease occurs in young adulthood and progresses with age [2]. According to the results of the Survey of Dental Diseases 2016, the percentage of persons in Japan with periodontal pockets of ≥4 mm increased with age [3]. Moreover, since 17.6% individuals aged 15-24 years and 32.4% individuals aged 25-34 years have periodontal pockets of ≥4 mm, periodontal disease is now known to be common among the elderly and the young adults. Among middle-aged individuals, the prevalence of periodontal disease is higher in women than in men. The decrease in estrogen levels after menopause leads to a decrease in bone density, which causes the progression of alveolar bone resorption, a symptom of periodontal disease [4]. Therefore, with age, women are at a greater risk of developing periodontal disease than men [4].

It is important to prevent periodontal disease because it is known to affect systemic diseases such as cardiovascular disease [5], diabetes mellitus [6, 7], and metabolic syndrome [8] and premature/low-birth-weight delivery [9, 10]. Previous studies assessing the causes and etiology of periodontal disease mainly focused on oral microbes and surrounding host environment [11, 12]. Previously reported environmental factors predisposing the host to periodontal disease include smoking habits, drinking habits [13], and dental hygiene [14].

In addition, the nutrients and food consumed have also been reported to be correlated with periodontal disease. Shariq Najeeb *et al.* suggested that decreased consumption of vitamins and minerals, particularly vitamin C, which has antioxidant activity, was highly associated with periodontal disease; however, vitamin C supplementation was not effective in treating periodontal disease [15]. Varun Kulkami *et al.* showed a similar intake of particular nutrients in patients with and without periodontal disease. However, they pointed to a possible correlation between consumed nutrients and periodontal disease status [16]. In addition, Yoshihara *et al.* showed a negative correlation between the intake of green and yellow vegetables and periodontal disease prevalence. However, it has also been reported that a high intake of cereals, nuts, sugar, sweets, and candy is positively correlated with the presence of periodontal disease [17]. As described above, most of the previous studies on the effect of diet/nutrient targeted elderly patients, with very few studies targeting the young adults.

Our foods are divided into relatively hard foods and soft foods based on physical properties. The hardness of food is regarded as the difficulty involved in chewing, hereafter referred to as dietary hardness. Dietary hardness has been assessed as an estimate of masticatory muscle activity for the habitual diet [28]. A positive correlation between dietary hardness and chewing muscle activity has also been reported in a previous study [18].

Therefore, we aimed at assessing the correlation between dietary habits and periodontal disease in young adult women. In addition, we assessed the correlation between the intake of hard foods and periodontal disease.

## 2. Materials and Methods

### 2.1. Participants

One hundred and twenty-seven female college students from the Tokyo Medical and Dental University and the Tokyo Healthcare University were recruited for the study between February 2014 to March 2015.

The inclusion criteria were not having caries teeth and missing teeth excluding third molars. On the other hand, the exclusion criteria were in the following; a smoker, taking gastrointestinal disease drugs, dry mouth, impaired oral movement and psychiatric illness.

After 6 smokers and 1 person taking gastrointestinal disease drugs were excluded, 120 subjects (mean age, 20.4 years) were finally enrolled in the study. One dentist with ≥5 years of experience in clinical trials assessed the subjects for periodontal disease. Two other evaluators practiced the chewing gum and gummy jelly tests in advance so that their results matched. The Ethical Committee of the Tokyo Medical and Dental University (#1002) and Tokyo Healthcare University (#25-8) approved the study protocols. All participants provided written informed consent before enrolment in the study, and experiments were performed following the guidelines in the Helsinki Declaration on the use of human subjects for research.

### 2.2. Dietary assessment

Nutrient/food intake was measured using the self-administered diet history questionnaire (DHQ). The validity and reliability of the DHQ have previously been confirmed [19, 20]. The DHQ includes questions concerning the frequency of consumption, the semi-quantitative portions of 151 food and beverage items consumed, general eating habits including skipping meals and eating irregularly, use of dietary supplements, and major cooking methods. An *ad hoc* of food composition in Japan was used to calculate the estimated dietary intake of energy, food group items, and nutrients [21]. The nutrient/food intake was converted to intake per 1,000 kcal of consumed energy. Since the total intake was expected to vary with body size, physical activity, and energy intake the nutrient/food intake was adjusted for total energy by using energy density (per 1,000 kcal) [22].

### 2.3. Estimation of dietary hardness

Dietary hardness was calculated using a previously described method [23]. Each food intake was multiplied by each masticatory muscle’s activity for the habitual diet. Subsequently, dietary hardness was obtained by dividing the total sum of the above value by the energy intake.

### 2.4. Periodontal disease variables

Each participant underwent assessments for periodontics according to the Community Periodontal Index (CPI) from code 0 to code 4 (code 0: health periodontal conditions; code 1: gingival bleeding on probing; code 2: calculus and bleeding; code 3: periodontal pocket 4–5 mm; and code 4: periodontal pocket ≥6 mm) [24, 25]. Then, the participants were assigned to two groups, namely, non-periodontal disease (non-PD) group for CPI codes 0-2 and periodontal disease (PD) group for CPI codes 3-4.

### 2.5. Anthropometry

Age and self-reported body heights were obtained from the DHQ. The weight and percentage of body fat were measured using a body composition meter (Inner scan 50, TANITA corporation, Japan). Body mass index (BMI) was calculated as body weight (kilograms) divided by the square of body height (meters).

### 2.6. Measuring masticatory performance by using color-changeable chewing gum and gummy jelly test

Mixing ability was measured using a color-changeable chewing gum (Masticatory Performance Evaluating Gum XYLITOL, Lotte Co., Ltd, Tokyo Japan). After mouth-rinsing with water for 15 seconds, the participants were then allowed to chew the provided gum by free chewing at a rate of one stroke per second with an electric metronome. The participants chewed the gum for 60 strokes. Changes in color were assessed using a colorimeter (CR-13, Konica-Minolta sensing, Tokyo, Japan), and then the mean values of L*, a*, and b* in the CIELAB color system were measured. Thereafter, the ΔE values were obtained using following the equation [26, 27].

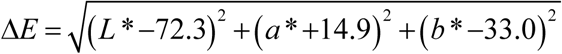

Comminution test was measured using a test gummy jelly (UHA Mikakuto Co., Ltd, Japan) and a reported scoring method [28]. The participants chewed the test jelly on their preferred chewing side for a total of 30 strokes. Subsequently, the chewed test jellies were evaluated on a scale of one to ten by using a visual scoring method [29]. Higher scores represented better comminution.

### 2.7. Health-related QoL

The general health-related QoL of the participants was evaluated by using the Health-related quality of life scale, Short Form-8 (SF-8) [30, 31, 32].

### 2.8. Statistical analyses

Age, height, weight, BMI, body fat percentage, nutrient/food intake, dietary hardness, masticatory performance, and health-related QoL were compared between the two groups by using Student’s *t*-test or Mann-Whitney U test. JMP Pro 11.0.0 (SAS) software was used for analyses. *P* < 0.05 was considered statistically significant. Except for physical status, dietary hardness, and QoL, all the other variables were log-transformed to achieve a normal distribution.

## 3. Results

### 3.1. Characteristics of participants

The mean age, body fat percentage, weight, height, and BMI of each group are shown in Table 1. The average age of the participants was 20.4 ± 1.1 years. Seventy-one subjects were assigned to the non-PD group, and 49 subjects were assigned to the PD group. There was no significant difference in age, height, weight, percentage body fat, and BMI between the two groups.

**Table 1.** The character of participants.

|  |  | Total |  | non-PD |  | PD |  | *p* |
| --- | --- | --- | --- | --- | --- | --- | --- | --- |
|  |  | Mean | SD | Mean | SD | Mean | SD |  |
| Age<sup>2)</sup> | yrs | 20.4 | 1.1 | 20.4 | 1.1 | 20.3 | 1.1 | 0.10 |
| Body height<sup>2)</sup> | cm | 158.9 | 5.8 | 158.4 | 5.3 | 159.0 | 6.3 | 0.91 |
| Body weight<sup>1)</sup> | kg | 51.2 | 6.9 | 51.2 | 7.4 | 51.1 | 6.2 | 0.89 |
| BMI<sup>2)</sup> | kg/m<sup>2</sup> | 20.3 | 2.4 | 20.4 | 2.6 | 20.2 | 2.0 | 0.78 |
| Body fat<sup>1)</sup> | % | 27.0 | 5.2 | 27.0 | 5.8 | 27.0 | 4.1 | 0.97 |

Total n=120; non-PD n=71; PD n=49
1) Student's t-test
2) Mann-Whitney U test

### 3.2. Intake of nutrients

Table 2 shows the results of the intake of nutrients. A comparison of energy intake and protein, fat, and carbohydrate energy ratio did not show a significant difference between the two groups. The intake of minerals such as potassium, calcium, magnesium, and iron was significantly lower in the PD group. Moreover, the intake of fat-soluble vitamins, including vitamin A, beta-carotene equivalence, vitamin E, and vitamin K, was significantly lower in the PD group. The intake of water-soluble vitamins, including pantothenic acid (vitamin B5), vitamin B6, and folic acid (vitamin B9), were also significantly lower in the PD group.

**Table 2.**
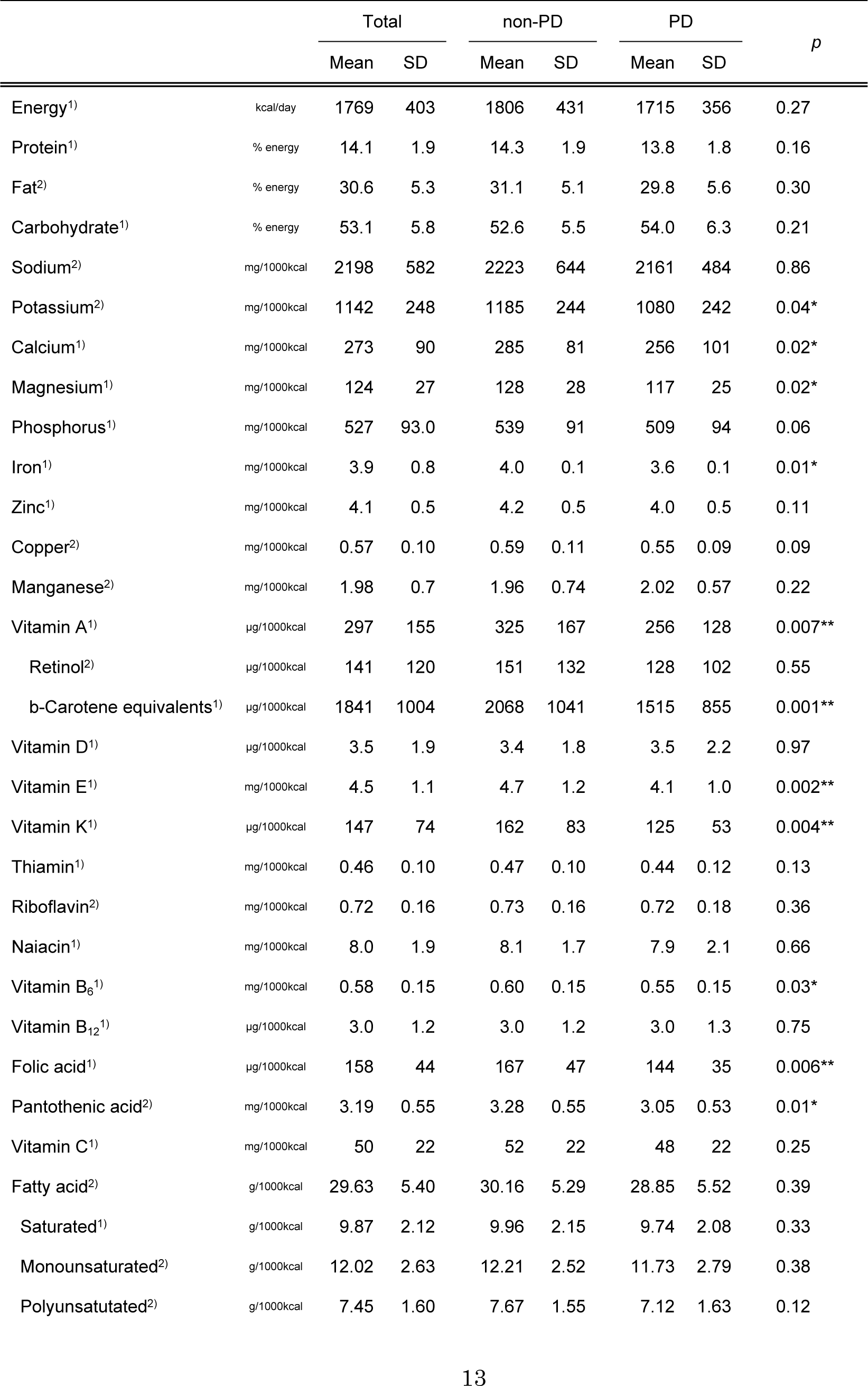
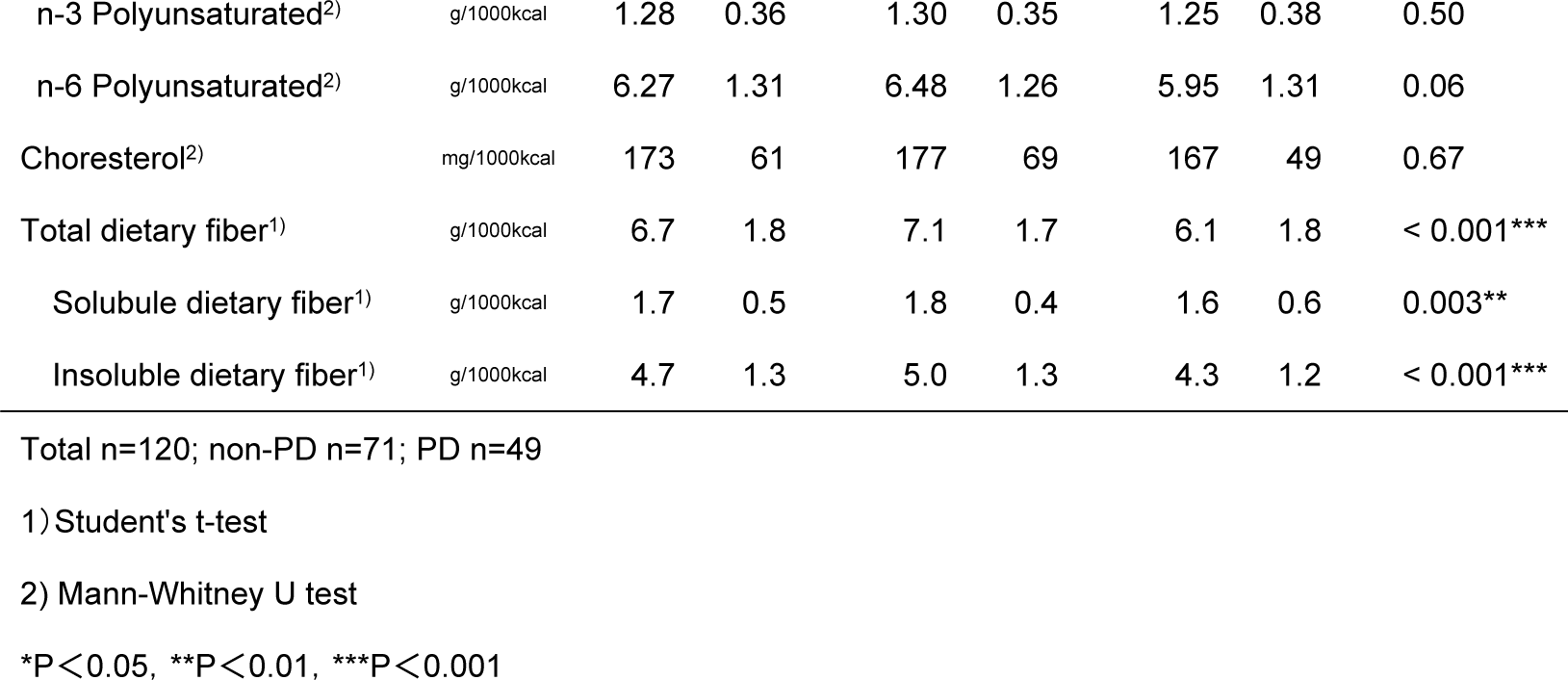
Nutrient intake in the PD and non-PD groups.

|  |  | Total |  | non-PD |  | PD |  | *p* |
| --- | --- | --- | --- | --- | --- | --- | --- | --- |
|  |  | Mean | SD | Mean | SD | Mean | SD |  |
| Energy<sup>1)</sup> | kcal/day | 1769 | 403 | 1806 | 431 | 1715 | 356 | 0.27 |
| Protein<sup>1)</sup> | % energy | 14.1 | 1.9 | 14.3 | 1.9 | 13.8 | 1.8 | 0.16 |
| Fat<sup>2)</sup> | % energy | 30.6 | 5.3 | 31.1 | 5.1 | 29.8 | 5.6 | 0.30 |
| Carbohydrate<sup>1)</sup> | % energy | 53.1 | 5.8 | 52.6 | 5.5 | 54.0 | 6.3 | 0.21 |
| Sodium<sup>2)</sup> | mg/1000kcal | 2198 | 582 | 2223 | 644 | 2161 | 484 | 0.86 |
| Potassium<sup>2)</sup> | mg/1000kcal | 1142 | 248 | 1185 | 244 | 1080 | 242 | 0.04* |
| Calcium<sup>1)</sup> | mg/1000kcal | 273 | 90 | 285 | 81 | 256 | 101 | 0.02* |
| Magnesium<sup>1)</sup> | mg/1000kcal | 124 | 27 | 128 | 28 | 117 | 25 | 0.02* |
| Phosphorus<sup>1)</sup> | mg/1000kcal | 527 | 93.0 | 539 | 91 | 509 | 94 | 0.06 |
| Iron<sup>1)</sup> | mg/1000kcal | 3.9 | 0.8 | 4.0 | 0.1 | 3.6 | 0.1 | 0.01* |
| Zinc<sup>1)</sup> | mg/1000kcal | 4.1 | 0.5 | 4.2 | 0.5 | 4.0 | 0.5 | 0.11 |
| Copper<sup>2)</sup> | mg/1000kcal | 0.57 | 0.10 | 0.59 | 0.11 | 0.55 | 0.09 | 0.09 |
| Manganese<sup>2)</sup> | mg/1000kcal | 1.98 | 0.7 | 1.96 | 0.74 | 2.02 | 0.57 | 0.22 |
| Vitamin A<sup>1)</sup> | µg/1000kcal | 297 | 155 | 325 | 167 | 256 | 128 | 0.007** |
| Retinol<sup>2)</sup> | µg/1000kcal | 141 | 120 | 151 | 132 | 128 | 102 | 0.55 |
| b-Carotene equivalents<sup>1)</sup> | µg/1000kcal | 1841 | 1004 | 2068 | 1041 | 1515 | 855 | 0.001** |
| Vitamin D<sup>1)</sup> | µg/1000kcal | 3.5 | 1.9 | 3.4 | 1.8 | 3.5 | 2.2 | 0.97 |
| Vitamin E<sup>1)</sup> | mg/1000kcal | 4.5 | 1.1 | 4.7 | 1.2 | 4.1 | 1.0 | 0.002** |
| Vitamin K<sup>1)</sup> | µg/1000kcal | 147 | 74 | 162 | 83 | 125 | 53 | 0.004** |
| Thiamin<sup>1)</sup> | mg/1000kcal | 0.46 | 0.10 | 0.47 | 0.10 | 0.44 | 0.12 | 0.13 |
| Riboflavin<sup>2)</sup> | mg/1000kcal | 0.72 | 0.16 | 0.73 | 0.16 | 0.72 | 0.18 | 0.36 |
| Niacin<sup>1)</sup> | mg/1000kcal | 8.0 | 1.9 | 8.1 | 1.7 | 7.9 | 2.1 | 0.66 |
| Vitamin B<sub>6</sub><sup>1)</sup> | mg/1000kcal | 0.58 | 0.15 | 0.60 | 0.15 | 0.55 | 0.15 | 0.03* |
| Vitamin B<sub>12</sub><sup>1)</sup> | µg/1000kcal | 3.0 | 1.2 | 3.0 | 1.2 | 3.0 | 1.3 | 0.75 |
| Folic acid<sup>1)</sup> | µg/1000kcal | 158 | 44 | 167 | 47 | 144 | 35 | 0.006** |
| Pantothenic acid<sup>2)</sup> | mg/1000kcal | 3.19 | 0.55 | 3.28 | 0.55 | 3.05 | 0.53 | 0.01* |
| Vitamin C<sup>1)</sup> | mg/1000kcal | 50 | 22 | 52 | 22 | 48 | 22 | 0.25 |
| Fatty acid<sup>2)</sup> | g/1000kcal | 29.63 | 5.40 | 30.16 | 5.29 | 28.85 | 5.52 | 0.39 |
| Saturated<sup>1)</sup> | g/1000kcal | 9.87 | 2.12 | 9.96 | 2.15 | 9.74 | 2.08 | 0.33 |
| Monounsaturated<sup>2)</sup> | g/1000kcal | 12.02 | 2.63 | 12.21 | 2.52 | 11.73 | 2.79 | 0.38 |
| Polyunsaturated<sup>2)</sup> | g/1000kcal | 7.45 | 1.60 | 7.67 | 1.55 | 7.12 | 1.63 | 0.12 |
| n-3 Polyunsaturated<sup>2)</sup> | g/1000kcal | 1.28 | 0.36 | 1.30 | 0.35 | 1.25 | 0.38 | 0.50 |
| n-6 Polyunsaturated<sup>2)</sup> | g/1000kcal | 6.27 | 1.31 | 6.48 | 1.26 | 5.95 | 1.31 | 0.06 |
| Cholesterol<sup>2)</sup> | mg/1000kcal | 173 | 61 | 177 | 69 | 167 | 49 | 0.67 |
| Total dietary fiber<sup>1)</sup> | g/1000kcal | 6.7 | 1.8 | 7.1 | 1.7 | 6.1 | 1.8 | < 0.001*** |
| Soluble dietary fiber<sup>1)</sup> | g/1000kcal | 1.7 | 0.5 | 1.8 | 0.4 | 1.6 | 0.6 | 0.003** |
| Insoluble dietary fiber<sup>1)</sup> | g/1000kcal | 4.7 | 1.3 | 5.0 | 1.3 | 4.3 | 1.2 | < 0.001*** |

Total n=120; non-PD n=71; PD n=49
1) Student's t-test
2) Mann-Whitney U test
\*P<0.05, \*\*P<0.01, \*\*\*P<0.001

### 3.3. Intake of different food groups

The results of food intake per 1000 kcal of energy for each group are shown in Table 3. The intake of green and yellow vegetables and other vegetables was significantly lower in the PD group than in the non-PD group. Although there was no significant difference in the intake of seeds, nuts, and beans, their intake was lower in the PD group. In addition, the intake of sugar, animal fats, and beverages was higher in the PD group. When green and yellow vegetables and other vegetables were analyzed (Table 4), the intake of carrots (among green and yellow vegetables) and the intake of cabbage (among other vegetables) were significantly lower in the PD group than in the non-PD group. Furthermore, although there was no significant difference, the intake of broccoli, as a green and yellow vegetable, and that of cucumber, eggplant, and lotus root, as a part of the other vegetables, were lower in the PD group.

**Table 3.** Food intake per 1000 kcal in the PD and non-PD groups. g/1,000kcal

|  | Total |  | non-PD |  | PD |  | *p* |
| --- | --- | --- | --- | --- | --- | --- | --- |
|  | Mean | SD | Mean | SD | Mean | SD |  |
| Cereals<sup>2)</sup> | 204.3 | 58.2 | 197.9 | 59.2 | 213.5 | 56.2 | 0.19 |
| Nuts and seeds<sup>2)</sup> | 1.1 | 2.5 | 1.2 | 2.6 | 0.9 | 2.2 | 0.09 |
| Potatoes<sup>2)</sup> | 13.9 | 8.4 | 14.8 | 8.4 | 12.7 | 8.4 | 0.06 |
| Sugars<sup>2)</sup> | 6.2 | 3.0 | 6.1 | 2.6 | 6.3 | 3.6 | 0.86 |
| Confectioneries<sup>2)</sup> | 51.4 | 25.1 | 51.3 | 25.3 | 51.6 | 25.1 | 0.89 |
| Animal fat<sup>2)</sup> | 0.5 | 0.7 | 0.4 | 0.5 | 0.6 | 0.8 | 0.20 |
| Vegetable fats and oil<sup>2)</sup> | 12.4 | 8.5 | 12.4 | 8.3 | 12.4 | 9.0 | 0.59 |
| Pulses<sup>1)</sup> | 38.2 | 41.1 | 42.9 | 48.4 | 31.5 | 26.2 | 0.17 |
| Fruits<sup>2)</sup> | 45.6 | 38.1 | 46.7 | 41.6 | 43.9 | 32.6 | 0.93 |
| Green and yellow vegetables<sup>1)</sup> | 61.4 | 35.9 | 68.5 | 38.5 | 51.2 | 29.4 | 0.01* |
| Other vegetables<sup>1)</sup> | 61.6 | 34.4 | 67.3 | 37.7 | 53.3 | 27.5 | 0.02* |
| Mushrooms<sup>2)</sup> | 7.3 | 7 | 7.3 | 6.5 | 7.4 | 8.7 | 0.21 |
| Seaweeds<sup>2)</sup> | 7.4 | 7.3 | 7.6 | 6.7 | 7.0 | 8.2 | 0.57 |
| Seasonings<sup>2)</sup> | 8.5 | 6.0 | 8.2 | 4.9 | 8.8 | 7.3 | 0.91 |
| Alcoholic beverages<sup>2)</sup> | 25.3 | 50.8 | 25.9 | 52.7 | 25.6 | 48.5 | 0.58 |
| Beverages<sup>2)</sup> | 383.7 | 241.4 | 357.5 | 244.0 | 421.7 | 234.8 | 0.09 |
| Fish<sup>1)</sup> | 28.9 | 15.2 | 28.4 | 15.3 | 29.5 | 15.2 | 0.84 |
| Meat<sup>1)</sup> | 43.3 | 18.9 | 44.4 | 17.1 | 41.7 | 21.3 | 0.17 |
| Eggs<sup>2)</sup> | 19.3 | 13.5 | 20.2 | 15.1 | 18.1 | 10.9 | 0.76 |
| Milk produces<sup>2)</sup> | 69.3 | 61.6 | 71.4 | 54.9 | 66.2 | 70.8 | 0.28 |

Total n=120; non-PD n=71; PD n=49
1) Student's t-test
2) Mann-Whitney U test
\*P<0.05

**Table 4.** Intake of green/yellow and other vegetables in the PD and non-PD groups. g/1,000kcal

|  | Total |  | non-PD |  | PD |  | *p* |
| --- | --- | --- | --- | --- | --- | --- | --- |
|  | Mean | SD | Mean | SD | Mean | SD |  |
| Green and yellow vegetables |  |  |  |  |  |  |  |
| Carrots<sup>2)</sup> | 8.1 | 5.9 | 9.1 | 5.9 | 6.7 | 5.8 | 0.02* |
| Pumpkins<sup>2)</sup> | 3.8 | 5.1 | 4.1 | 5.8 | 3.5 | 4.0 | 0.94 |
| Tomatoes<sup>1)</sup> | 13.0 | 12.5 | 14.5 | 13.6 | 10.9 | 10.3 | 0.13 |
| Sweet peppers<sup>2)</sup> | 2.9 | 3.3 | 3.1 | 3.6 | 2.6 | 2.8 | 0.36 |
| Broccoli<sup>2)</sup> | 6.6 | 9.3 | 7.7 | 10.2 | 5.1 | 7.8 | 0.06 |
| Green leafy vegetables<sup>2)</sup> | 17.7 | 18.5 | 20.8 | 22.3 | 13.3 | 9.5 | 0.15 |
| Other vegetables |  |  |  |  |  |  |  |
| Cabbage<sup>1)</sup> | 14.7 | 14.9 | 17.1 | 16.6 | 11.2 | 11.3 | 0.01* |
| Cucumbers<sup>1)</sup> | 4.8 | 5.7 | 5.5 | 5.5 | 3.8 | 4.3 | 0.05 |
| Lettuce<sup>2)</sup> | 6.3 | 5.8 | 7.3 | 6.5 | 5.0 | 4.2 | 0.13 |
| Chinese cabbage<sup>2)</sup> | 4.4 | 7.4 | 4.4 | 7.5 | 4.4 | 7.2 | 0.73 |
| Bean sprouts<sup>2)</sup> | 5.4 | 7.1 | 5.7 | 7.0 | 5.0 | 7.2 | 0.60 |
| Radishes<sup>2)</sup> | 6.4 | 6.0 | 7.0 | 6.5 | 5.6 | 5.2 | 0.43 |
| Onions<sup>1)</sup> | 10.9 | 9.5 | 11.3 | 10.2 | 10.3 | 8.6 | 0.59 |
| Cauliflower<sup>2)</sup> | 0.3 | 0.9 | 0.4 | 1.1 | 0.2 | 0.6 | 0.25 |
| Eggplants<sup>2)</sup> | 3.4 | 5.4 | 3.6 | 4.0 | 3.1 | 7.1 | 0.06 |
| Burdock<sup>2)</sup> | 1.7 | 2.3 | 2.0 | 2.8 | 1.2 | 1.1 | 0.12 |
| Lotus root<sup>2)</sup> | 0.8 | 1.1 | 0.9 | 1.3 | 0.7 | 0.9 | 0.06 |

Total n=120; non-PD n=71; PD n=49
1) Student's t-test
2) Mann-Whitney U test
\*P<0.05

### 3.4. Dietary hardness

Masticatory muscle activity was analyzed as a measurement of dietary hardness (Table 5). The results showed that the intake of hard foods was significantly lower in the PD group when compared with the non-PD.

**Table 5.** Hard food intake in the PD and non-PD groups. m·V/1000kcal

|  | Total |  | non-PD |  | PD |  | *p* |
| --- | --- | --- | --- | --- | --- | --- | --- |
|  | Mean | SD | Mean | SD | Mean | SD |  |
| Intake of hard foods<sup>1)</sup> | 204 | 30.4 | 208 | 31 | 197 | 30 | 0.046* |

Total n=120; non-PD n=71; PD n=49
1) Student's t-test
\*P<0.05

### 3.5. Masticatory performance and health-related QoL

Masticatory performance and QoL are shown in Table 6. There was no significant difference in mixing ability, comminution ability, and QoL between the two groups.

**Table 6.**
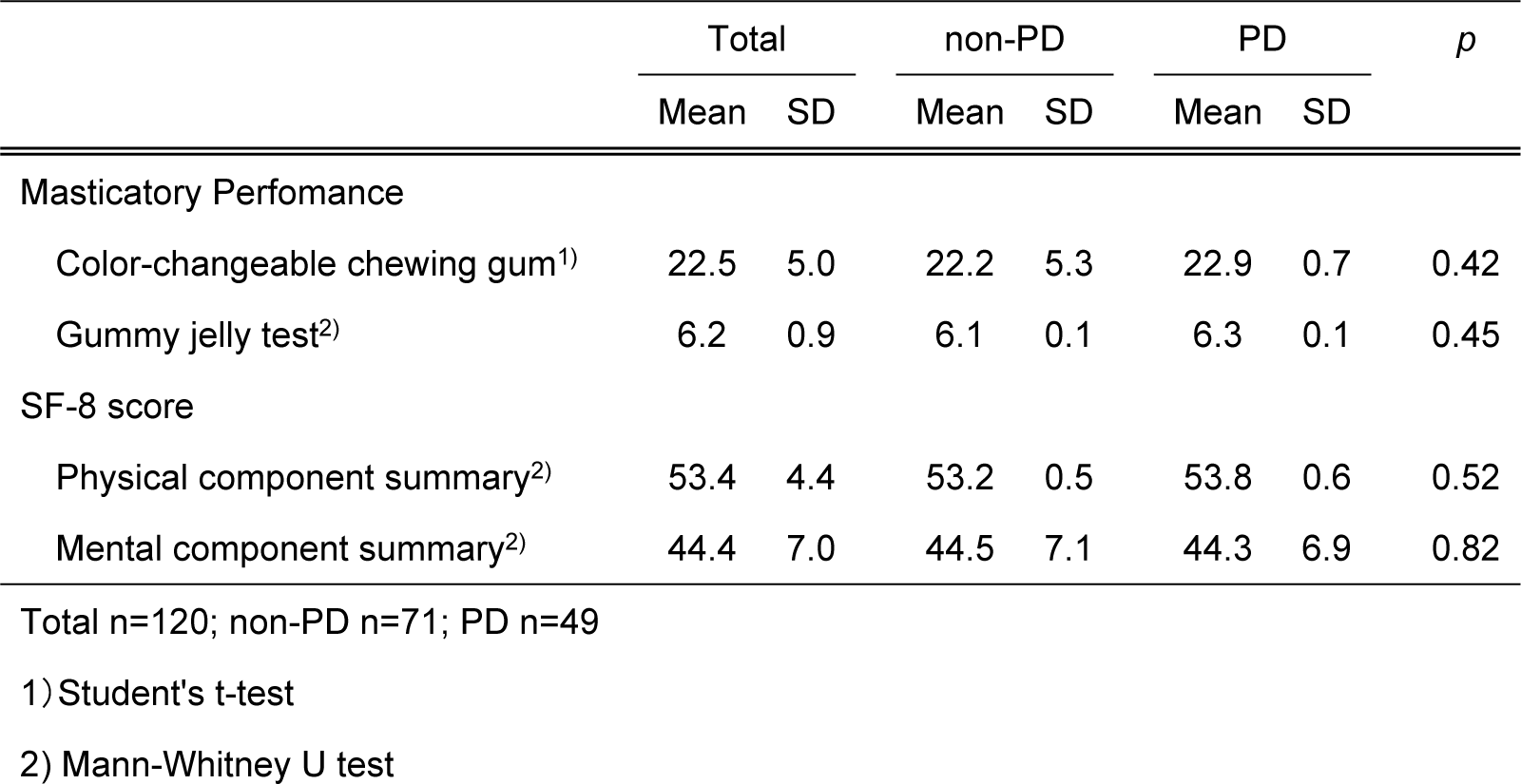
Masticatory performance and health related QoL.

## 4. Discussion

In this study, the two groups showed significant differences in the intake of several nutrients.

In previous studies, subjects with periodontal disease significantly consumed less potassium and iron. In the present study, the intake of potassium and iron in the PD group was significantly lower than that in the non-PD group, which was consistent with that reported in previous studies. Magnesium is known to be involved in bone and tooth formation. It was reported that in 40-year-old individuals, serum magnesium concentration is associated with the risk of periodontal disease [33]. The intake of magnesium was significantly lower in the PD group in the present study. We, therefore, think that there is a significant positive correlation between the presence of periodontal disease and low intake of magnesium in not only the elderly but also in young adults. Both animal and human studies have reported that a lack of calcium intake leads to alveolar bone resorption [34, 35] and a decrease in bone density [36], indicating that insufficient calcium intake might cause the progression of periodontal disease. In addition, it was shown that the risk of periodontal disease associated with calcium intake was observed in both men and women aged between 20 and 39 years (odds ratio; 1.99 and 1.84), only in men aged between 40 and 59 years, and in neither men nor women aged ≥60 years [37]. In the present study, the intake of calcium was significantly lower in the PD group. Therefore, we think that low calcium intake is more strongly correlated with the presence of periodontal disease in younger adults than in older adults.

In the present study, the intake of vitamin C was lower, although not significantly, in the PD group, while the intake of vitamins A, K, and E and beta-carotene were significantly lower. Among the elderly and smokers, the prevalence of periodontal disease was reported to be associated with vitamin C intake. There was a weak but significant dose-response increase in the risk of periodontal disease in subjects who had lower vitamin C intake [38]. Vitamin C is known to suppress oxidative stress through its antioxidant activity [39]. In addition, vitamins A and K [40] and vitamin E and beta-carotene [41] have antioxidant activity. In previous studies, oxidative stress caused by reactive oxygen species in the tissues surrounding the gingiva affects the onset of periodontal disease [42]. Therefore, it was suggested that the antioxidant effect of several vitamins contributed to the absence of periodontal disease in the non-PD group. Low intake of folic acid is known to be related to increased bleeding during probing [43, 44], which supports our findings that the intake of folic acid was significantly lower in the PD group. In the present study, the intake of pantothenic acid and vitamin B6, which are not known to have any correlation with periodontal disease, was significantly lower in the PD group. Previous reports have shown that decreased vitamin B complex intake reduced the efficiency of access flap surgery for periodontal disease treatment [45]. However, the effect of each specific vitamin B in the complex on periodontal disease treatment has not been shown yet. Based on our findings, it was suggested that there is a positive correlation between the intake of pantothenic acid and vitamin B6, among the vitamin B complex, and the absence of periodontal disease.

In a previous study, there was a significant positive correlation between the intake of whole grains and the absence of periodontal disease because whole grains contain dietary fiber [46]. Green and yellow vegetables and other vegetables like whole grains contain soluble and insoluble dietary fibers. Therefore, in this study, since the consumption of green/yellow vegetables and other vegetables was higher in the non-PD group than in the PD group, we think that dietary fibers play a protective role against the development of periodontal disease.

It was reported that the risk for periodontal disease among elderly persons with low intake of docosahexaenoic acid is approximately 1.5 times that among those with high intake [47]. However, in the present study, there was no significant difference in the intake of fatty acids between the two groups. N-3 polyunsaturated fatty acids such as docosahexaenoic acid and eicosapentaenoic acid exert anti-inflammatory effects by blocking intracellular signaling systems required for producing inflammatory cytokines such as IL-1, IL-6, and TNF-α [47]. Therefore, the immunological differences between younger adults and the elderly might have caused the difference in the findings between the present study and previous studies.

This study showed that the PD group significantly consumed foods with lower dietary hardness when compared with the non-PD group. Yanagisawa, *et al*. classified foods into 10 ranks according to chewing muscle activity [18]. In the study, carrots and cabbage were ranked 8^th^ out of 10 (>1400 μV· sec) and carrots, cabbage, and lotus root were ranked 6^th^ out of 10 (>1000 μV·sec). We found that the non-PD group routinely consumed hard dietary foods when compared with the PD group. The quantity of saliva secreted has been reported to increase proportionally with an increase in chew number, chewing time, or chewing muscle activity [48]. Saliva washes away residual food particles and bacteria in the mouth. Saliva also contains lysozyme, amylase, peroxidase, and the proteins lactoferrin and IgA which possess antibacterial activity [49, 50]. Therefore, we think that eating foods with high dietary hardness (as seen in the non-PD group) protects against periodontal disease through oral washing and antibacterial activity resulting from increased saliva secretion.

In the present study, the participants in the PD group accounted for 40.8% of the total participants. The 2016 Survey of Dental Diseases reported that women aged 20 to 24 with periodontal disease accounted for 16.7% of the total population [3]. Thus, the high prevalence of periodontal disease observed in our study population is likely because we did not comprehensively diagnose periodontal disease by using multiple diagnostic methods. Our diagnoses were based only on the CPI, which might have resulted in the observed high prevalence.

Presently, dietary or nutritional studies assessing how diet/nutrition can be helpful in the prevention or treatment of periodontal disease are lacking. Therefore, more studies are required to elucidate the connection between periodontal disease and diet/nutrition. In addition, longitudinal studies on dietary habits that effectively prevented or suppressed the progression of periodontal disease are warranted.

## 5. Conclusion

Under the limited conditions of the present study, the intake of minerals (potassium, calcium, magnesium, and iron), fat-soluble vitamins (vitamin A, beta-carotene, vitamin E, and vitamin K), water-soluble vitamins (pantothenic acid, vitamin B6, and folic acid), and dietary fibers was significantly lower in young adult women with periodontal disease than in young adult women without periodontal disease.

## Acknowledgments

The authors thank all the participants from the Tokyo Medical and Dental University and Tokyo Healthcare University.

